# Efficient isolation of protoplasts from rice calli with pause points and its application in transient gene expression and genome editing assays

**DOI:** 10.1101/2020.09.01.278176

**Authors:** Snigdha Poddar, Jaclyn Tanaka, Jamie H. D. Cate, Brian Staskawicz, Myeong-Je Cho

## Abstract

An efficient *in vivo* transient transfection system using protoplasts is an important tool to study gene expression, metabolic pathways, and multiple mutagenesis parameters in plants. Although rice protoplasts can be isolated from germinated seedlings or cell suspension culture, preparation of those donor tissues can be inefficient, time consuming, and laborious. Additionally, the lengthy process of protoplast isolation and transfection needs to be completed in a single day. Here we report a protocol for isolation of protoplasts directly from rice calli, without using seedlings or suspension culture. The method is developed to employ discretionary pause points during protoplast isolation and prior to transfection. Protoplasts maintained within a sucrose cushion partway through isolation, for completion on a subsequent day, per the first pause point, are referred to as S protoplasts. Fully isolated protoplasts maintained in MMG solution for transfection on a subsequent day, per the second pause point, are referred to as M protoplasts. Both S and M protoplasts, 1 day after initiation of protoplast isolation, had minimal loss of viability and transfection efficiency compared to protoplasts 0 days after isolation. S protoplast viability decreases at a lower rate over time than that of M protoplasts and can be used with added flexibility for transient transfection assays and time-course experiments. The protoplasts produced by this method are competent for transfection of both plasmids and ribonucleoproteins (RNPs). Cas9 RNPs were used to demonstrate the utility of these protoplasts to assay genome editing *in vivo*. The current study describes a highly effective and accessible method to isolate protoplasts from callus tissue induced from rice seeds. This method utilizes donor materials that are resource-efficient and easy to propagate, permits convenience via pause points, and allows for flexible transfection days after protoplast isolation. It provides an advantageous and useful platform for a variety of *in vivo* transient transfection studies in rice.

## Background

Rice (*Oryza sativa* L.) is a vital crop that provides staple calories for approximately half of the global population and is a model organism for basic research of monocotyledon plant biology [1, 2]. Amidst rapid population growth, climate change, and threats posed by pests and pathogens, the need to address food security via improved agricultural output is high. To meet these challenges, it is important to advance basic scientific understanding of plant processes, molecular machinery, and genetics. Concurrently, those advances can be applied and developed via biotechnological efforts to improve plants for increased yield, new genetic diversity, insect resistance, disease resistance, drought tolerance, herbicide tolerance and other agronomically important traits [3].

Much of this work, particularly early stage experiments, can be hastened via robust protoplast systems. The delivery of DNA or RNPs into plant tissue for biological assays is impeded by the presence of a rigid cell wall surrounding each cell. Enzymatic digestion of the cell walls followed by a purification process yields membrane-bound protoplasts [4]. These cells are useful and versatile gene expression systems competent for transfection of exogenous genetic material. Other experimental platforms exist in plants, such as heterologous expression in plants like onion or tobacco [5,6] and stable and transient transformation by *Agrobacteria* [7,8] or particle bombardment [9,10]. However, heterologous expression systems can be linked to a caveat of aberrant characteristics [11], and stable transformation requires significant resources and can be superfluous for some applications.

Protoplast studies are uniquely suited for facile, rapid, and high throughput *in vivo* assays to examine gene expression as well as to evaluate genome editing efficacy. The advent of targeted plant genome editing, mediated by various sequence-specific nucleases, is a powerful biotechnological development that has hastened plant gene function studies and crop development [12]. The CRISPR/Cas9 system, in particular, has provided significant utility due to its simplicity and versatility [3,13]. A critical factor driving editing efficiency is the guide RNA (gRNA) sequence that guides specific Cas9 cleavage of genomic DNA. Because generating stable genome-edited plants is complex and labor intensive, it is beneficial to first determine the most effective gRNAs *in vivo* as well as identify the range of mutations made through a simple and rapid protoplast pipeline.

Generally, protoplasts are isolated from leaves or germinated seedlings for transient transfection in several plant species [14,15,16,17,18]. Rice protoplasts can be isolated from cell suspension culture [19,20] as well as seedlings [21,22,23]. While effective, these methods can be time consuming and laborious. Isolation from seedlings requires 80-120 fresh seedlings per protoplast preparation, which can deplete seed pools quickly. Meticulous manual slicing of the plant material into small strips is also a critical step in the protocol. The blade must be changed regularly to ensure clean cuts, as any bruising of the leaf tissue leads to a lower yield of healthy protoplasts. Meanwhile, establishment and maintenance of cell suspension culture requires experienced skill to select proper callus morphologies and are vulnerable to contamination [19,24,25]. Furthermore, transfections are performed immediately after isolation, raising an additional component of time sensitivity.

In the present study, we describe a highly efficient method to isolate rice protoplasts from callus tissue derived from dry seeds. The induction and proliferation of calli is straightforward, sustainable, and sterile. We analyze protoplast viability and transfection competence over time, utilize the method for a genome editing assay, and demonstrate that this method provides convenience via pause points during protoplast isolation and is permissive for transfection of protoplasts for multiple days after initial cell wall digestion of calli.

## Materials and methods

### Plant materials

Plants of two rice (*Oryza sativa* L.) cultivars, Nipponbarre and Kitaake, were grown in a greenhouse at 16/8-h photoperiod intervals (250-300 μmol m^-2^s^-1^), 27 °C and 22 °C, respectively.

### Reagents and solutions

Recipes for callus induction media, digestion solution, W5 solution, MMG solution, WI solution, and PEG-CaCl_2_ solution are listed in Table 1. All solutions are 0.2 μm filter sterilized.

**Table 1.** Compositions of medium/solutions used for protoplast isolation and transfection

| Medium/Solution Name | Compositions |
| --- | --- |
| OsCIM2 | 3.99 g/L Chu's N6 Basal Medium with Vitamins (C167, PhytoTechnology Laboratories, Lenexa, KS, USA), 30 g/L maltose, 0.1 g/L myo-inositol, 5 uM CuSO<sub>4</sub>, 0.3 g/L casein enzymatic hydrolysate, 2.5 mg/L 2,4-D, 0.2 mg/L BAP, 0.5 g/L L-Proline, 0.5 g/L L-Glutamine, pH 5.8, solidified with 3.5 g/L Phytigel (P8169; Sigma-Aldrich Corp., St Louis, MO, USA). Autoclaved. |
| Digestion solution | 10 mM MES pH 5.7, 0.6 M mannitol, 1.5% cellulase Onozuka R-10 (Yakult, Tokyo, Japan), 0.1% pectolyase (or 0.75% macerozyme R-10) (Yakult, Tokyo, Japan), 10 mM CaCl<sub>2</sub>, 4 mM 2-mercaptoethanol, 0.1% bovine serum albumin.<br>*Special instructions:* MES, mannitol, H<sub>2</sub>O, cellulase R10, and pectolyase were stirred and incubated at 55°C for 10 minutes. The solution was cooled to room temperature, and CaCl<sub>2</sub>, 2-mercaptoethanol, and bovine serum albumin were added in and gently mixed. |
| W5 solution | 2 mM MES pH 5.7, 154 mM NaCl, 5 mM KCl, 125 mM CaCl<sub>2</sub> |
| MMG solution | 4 mM MES pH 5.7, 0.6 M mannitol, 15 mM MgCl<sub>2</sub> |
| WI solution | 4 mM MES pH 5.7, 0.4 M mannitol, 4 mM KCl |
| PEG-CaCl<sub>2</sub> solution | 0.4 M mannitol, 100 mM CaCl<sub>2</sub>, 40% (wt/vol) PEG4000 (81240; Sigma-Aldrich Corp., St Louis, MO, USA) |

### Callus induction and subculture

Mature seeds of rice were used to induce callus tissue. Briefly, dehulled mature seeds were surface sterilized for 15-20 minutes in 20% (v/v) bleach (5.25% sodium hypochlorite) plus one drop of Tween 20 followed by three washes in sterile water and placed on OsCIM2 callus induction medium (Table 1). After 7-14 days, the coleoptiles and endosperm tissues were removed from the mature seeds and the translucent, pale-yellow nodular calli were transferred onto fresh OsCIM2 every 3 to 4 weeks for subculture.

### Protoplast isolation

All exposed steps were performed under sterile conditions within a laminar flow hood. After enzymatic digestion of tissue, all pipetting of protoplasts was performed with sterile 1 mL tips with the top 0.25 cm removed. Five to six grams of compact, nodular callus tissue were collected from subcultured OsCIM2 plates and gently crumbled using the edge of a metal spatula or scalpel in a deep 25×100 mm petri dish with 15 mL of digestion solution. The petri dish was incubated in the dark in a room temperature shaker at 70 rpm for 3 hours until the digestion solution appeared milky.

The protoplast-filled digestion solution was first filtered through a Falcon 100 μm nylon cell strainer (352360; BD Biosciences, San Jose, CA, USA) in a sterile petri dish and then through a Falcon 40 μm nylon cell strainer (352340; BD Biosciences). The protoplast solution was transferred to a 50 mL conical tube and centrifuged for 5 minutes at room temperature at 150x*g*. The supernatant was discarded, and the protoplast pellet was gently resuspended in 8 mL W5 solution. Separately a fresh 50 mL conical tube with 10 mL of 0.55 M sucrose was prepared. The cell suspension was gently pipetted onto the sucrose cushion such that the cell suspension floated on top, then centrifuged at 1000x*g* without deceleration for 5 minutes.

At this stage, the isolation process could be paused until subsequent days, or continued immediately.

The intermediate cloudy phase, containing live protoplasts, was pipette extracted and mixed with 10 mL W5 solution in fresh tubes. The suspension was centrifuged for 5 minutes at room temperature at 150x*g*. The supernatant was removed, and the pellet gently resuspended in 5 mL of MMG solution. The suspension was once again centrifuged for 5 minutes at room temperature at 150x*g*. The protoplast pellet was resuspended in 4 mL of MMG, or enough to bring the final cell concentration to 2.5×10^6^ cells/mL as calculated by microscopy on a hemocytometer.

### Protoplast viability

One μL of 1% Evans blue (E2129; Sigma-Aldrich Corp., St Louis, MO, USA) was added to 25 μL protoplast suspension. The protoplasts were viewed on a hemocytometer under a light microscope. Live protoplasts, which remained unstained, were counted and total live protoplasts per milliliter were calculated. Dead protoplasts and debris were stained blue.

### Protoplast transfection

PEG-mediated transfection was performed, guided by previously published methods [22,26] with modifications. In a sterile 1.5 mL tube, 10 μg of 250 ng/μL plasmid DNA pAct1IsGFP [27] were added to 200 μL of protoplasts suspension (5×10^5^ total cells), gently flicked and inverted to mix thoroughly, and incubated at room temperature in the dark for 5 minutes. Two hundred forty μL of PEG-CaCl_2_ solution were added, and the tube inverted gently several times until fully mixed. This was further incubated at room temperature in the dark for 20 minutes. After incubation, 800 μL of W5 solution were added to stop the reaction, inverted gently several times until fully mixed, and centrifuged at 200x*g* for 5 minutes. The supernatant was carefully pipetted for removal, reserving the protoplast pellet. The protoplast pellet was resuspended with gentle inversions and minimal pipetting in 1 mL WI solution and transferred into a 12-well tissue culture plate. The plate edges were sealed with parafilm and incubated in the dark at 26 °C for 48 hours until they were utilized for light microscopy to measure protoplast viability on a hemocytometer and GFP fluorescence using a Zeiss Axio Imager (Carl Zeiss Microscopy LLC, White Plains, NY) and a Leica M165 fluorescence microscope (Leica Microsystems Inc., Buffalo Grove, IL).

### Rice protoplast genome editing and amplicon next generation sequencing analysis

Protoplasts were transfected with Cas9 RNPs based on a previous study, with modifications [28]. A 1:1 ratio of tracrRNA and target specific crRNA (Integrated DNA Technologies, Coralville, IA) were annealed to form gRNA. Ten μg Cas9 protein (Macrolab, University of California, Berkeley, CA) and 10 ug gRNA were incubated at 37 °C for 20 minutes in a total 25 μL to assemble the Cas9 RNPs. Protoplast transfection was performed, as described above, using 25 μL RNPs instead of plasmid DNA. Forty-eight hours post-transfection, the protoplasts were harvested for CTAB-chloroform genomic DNA extraction. To determine mutation rates by amplicon sequencing, PCR was performed with target specific primers, amplifying approximately 300 bp around the cut site using Q5 High-Fidelity (New England Biolabs, Ipswich, MA) polymerase. Primers contained a 5’-stub compatible with Illumina NGS library preparation. PCR products were ligated to Illumina TruSeq adaptors and purified. Libraries were prepared using a NEBNext kit (Illumina) according to the manufacturer’s guidelines. Samples were deep sequenced on an Illumina MiSeq at 300 bp paired-end reads to a depth of approximately 10,000 reads per sample. Cortado (https://github.com/staciawyman/cortado) was used to analyze editing outcomes. Briefly, reads were adapter trimmed and then merged using overlap to single reads. These joined reads were then aligned to the target reference sequence. Editing rates are calculated by counting any reads with an insertion or deletion overlapping the cut site or occurring within a 3 base pair window on either side of the cut site. SNPs occurring within the window around the cut site are not counted. Total edited reads are then divided by the total number of aligned reads to get percent edited.

## Results and Discussion

### A sustainable protoplast isolation method with optional pause points

Existing methods for protoplast isolation from rice using germinated seedlings and suspension cultures are valuable and well described [19–25]. However, they can consume seeds at a high rate or require the technical and labor-intensive know-how of maintaining a suspension culture. Furthermore, the methods require an uninterrupted lengthy workflow from donor tissue digestion all the way through transfection performed on the isolated protoplasts.

Here, techniques are outlined for a branched method with built-in optional pause points that allow for consistent and efficient procurement of healthy protoplasts that may be used gradually, over the course of several days, for downstream transient assays. The donor tissue for the isolation of protoplasts are calli induced from seeds and regularly sub-cultured on solid OsCIM2 callus induction media (Table 1). Calli were also induced from immature embryos in this manner, with comparable outcomes. In general, callus tissue propagated for more than six months could be used in this method. As a result, the donor tissue becomes available at an exponential rate once initiated, abrogating the obstacle of donor material availability for this procedure.

The workflow for the protoplast isolation protocol produced from this study is portrayed in Fig. 1. We gathered 5 g of compact pale-yellow rice callus tissue (Fig. 2A) and used a scalpel or metal spatula to bring all of the pieces to roughly the same size. Careful slicing and razor exchanges were not needed. An enzymatic cocktail of 1.5% cellulase R10 and 0.1% pectolyase or 0.75% macerozyme R10 resulted in successful breakdown of rice callus tissue cell walls while maintaining healthy viable protoplasts (Figs. 2B, C). An additional step of vacuum infiltration of the digestion solution with the donor tissue, an approach utilized in other protocols [22,29], was unnecessary for our method, eliminating a common step, decreasing equipment load, and increasing simplicity. Rather, we could simply incubate the callus tissue with 15 mL digestion solution with gentle shaking at 70 rpm for 3 hours, less time than is required for seedling-derived cells. After digestion, protoplasts were isolated from spent tissue via filtration (Fig. 2D) and centrifugation through a 0.55 M sucrose cushion. A gentle overlay of the cell suspension onto the sucrose was found to be a crucial step for optimal yield. If the cells were handled crudely and dropped with a force that significantly broke the surface tension of the sucrose, the ultimate protoplast yield could be diminished. After centrifugation of the cell suspension through 0.55 M sucrose, healthy protoplasts separated from debris and accumulated to form a dense band of purified protoplasts at the W5 - sucrose interface (Fig. 2E), bringing the protocol to its first optional pause point.

**Figure 1.**
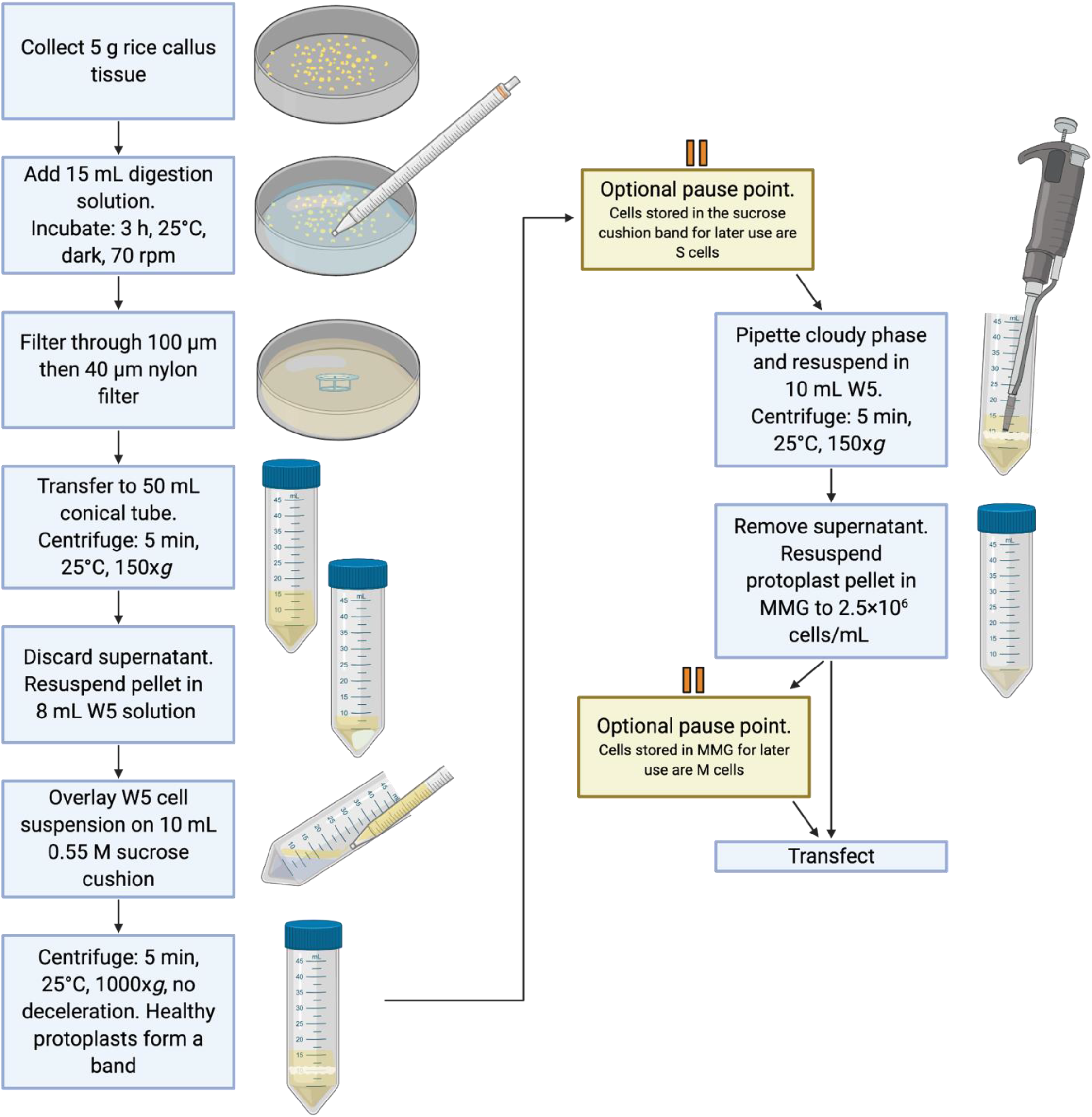
Graphical depiction of the protoplast isolation workflow.

**Figure 2.**
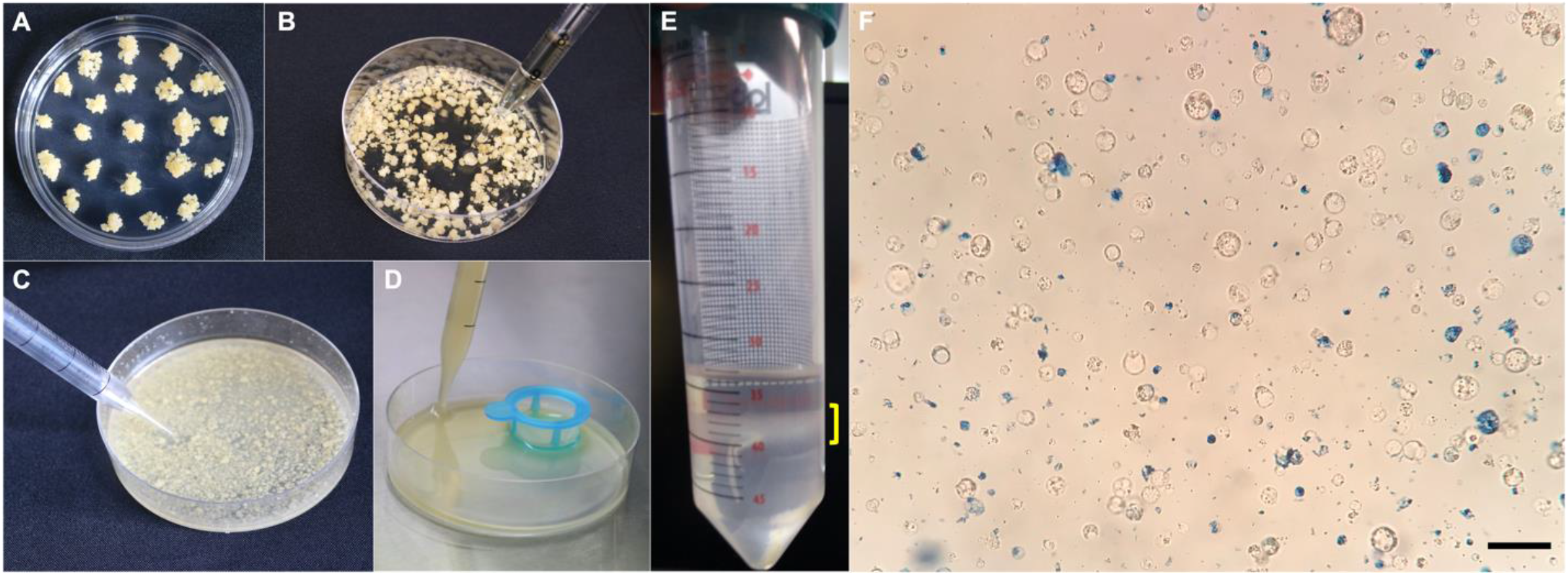
Isolation of protoplasts from rice calli induced from mature seeds. (A) Donor tissue for protoplast isolation were translucent, pale yellow, and nodular calli propagated on OsCIM2 medium. (B) To degrade the tissue cell walls, digestion solution was added. (C) A visual indication of successful enzymatic digestion was a milky appearance of the solution after three hours of gentle shaking. (D) Large particles and spent tissue were removed from the protoplast solution via cell strainer filtration. (E) Protoplasts formed a visible band, marked by a red bracket, after centrifugation through a 0.55 M sucrose cushion. (F) Protoplasts derived from rice calli. Healthy cells are round and colorless. Dead cells and debris are stained by Evans Blue. Bar = 50 μm

Here, the method could be paused for one or more days. The band of protoplasts could be left undisturbed at the interface for processing at a later time or handled immediately. The protoplasts produced from utilization of this pause point are termed “S protoplasts.”

To isolate protoplasts from the sucrose cushion, the entirety of the cloudy phase band containing the protoplasts was gently pipetted out. This was followed by final washing and centrifugation steps to maximize purity of the protoplasts and eliminate cellular debris. Finally, protoplasts were resuspended in MMG solution to a concentration of 2.5 x 10^6^ protoplasts/mL. Protoplasts could be transfected immediately, or the second optional pause point could be employed—storing the protoplasts in MMG solution, termed “M protoplasts,” until transfection at a later time.

Protoplasts isolated from the band on the same day as digestion of the donor tissue cell walls were referred to as “Sucrose Cushion Day 0/ MMG Day 0” (S/M0) protoplasts. Those isolated from the interface one, two, three, or 7 days after digestion were designated S1, S2, S3, or S7 protoplasts, respectively. S/M0 protoplasts, stored in MMG solution and utilized in experiments over the following one, two, three, or 7 days after digestion were labeled “MMG Day 1” (M1), M2, M3, or M7 protoplasts.

### Protoplast viability over time

To ensure the utility of this branched method for protoplast isolation, Evans blue staining was used to quantify viable protoplasts in all isolations from the sucrose cushion (S protoplasts) as well as protoplasts stored in MMG solution over time (M protoplasts) (Fig. 2F). Healthy intact protoplasts derived from this method are colorless, spherical, and resistant to staining. The viability assay indicated that S/M0 isolates yielded the greatest number of live protoplasts, with a gradual decrease with increasing age of the protoplast-containing sucrose cushion. S/M0 isolates contained approximately 2.5 times the number of protoplasts as S7 isolates. However, it is notable that the order of magnitude for the protoplast count remained unchanged between S/M0 and S7. We show that from 5 grams of rice callus donor tissue, this method yields, on average, 9.8×10^6^ live protoplasts if isolated on day 0 (S/M0) and 3.9×10^6^ live protoplasts if isolated on day 7 (S7) (Fig. 3A). This translates to approximately 20 transfection reactions with S/M0 protoplasts, and 8 transfection reactions with S7 protoplasts. To compare, 100-120 finely sliced rice seedlings are required to obtain approximately the same number of protoplasts as an S/M0 isolation by the current method [22], and which cannot be stored for future use.

**Figure 3.**
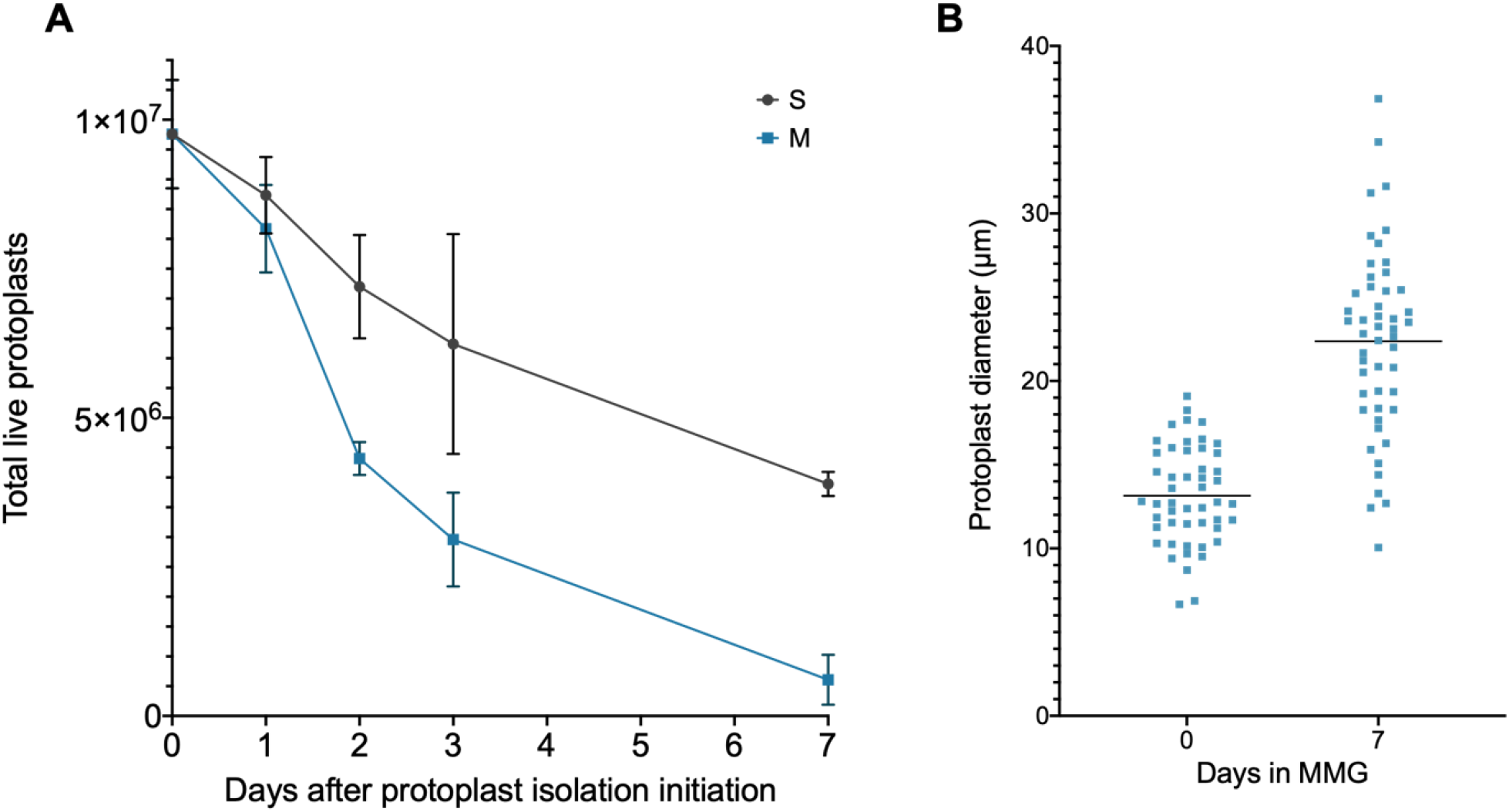
Protoplast viability and size over time. (A) Viability of protoplasts was measured by counting protoplasts unstained by Evans Blue dye on a hemocytometer. Total live protoplasts were calculated by first determining protoplast density, then multiplying by the total volume of protoplasts from the isolation. Counts were performed in triplicate. The means are plotted and error bars indicate standard deviation. (B) A random sampling of 50 protoplasts were measure for diameter at 0 and 7 days in MMG. Each measurement was plotted individually, and the means were indicated by a horizontal line.

Viability of M protoplasts was also tracked over time. Though viability decreases appreciably from day 2, the concentration of viable M1 protoplasts is comparable to S1 protoplasts and only mildly reduced from S/M0 protoplasts, making them an acceptable option for use in assays (Fig. 3A). It was also noted that protoplasts held in MMG solution for 7 days appeared approximately 1.7X larger (Fig. 3B). This may be attributed to cell growth or osmotic swelling. However, the larger protoplasts displayed a characteristically healthy spherical shape, unstained by Evans blue, suggesting that osmotic stress was not occurring.

### Transfection efficiency over time

Both quantity and quality of protoplasts are critical factors for downstream experiments. In existing methods, transfection is performed only on freshly isolated protoplasts. Here, transfection efficiency of both S and M protoplasts of different ages were assayed via PEG-mediated transfection of pAct1IsGFP-1, a GFP overexpression plasmid [27].

First, S and M transfection pools were imaged for GFP expression 1 and 2 days after transfection (Fig. 4). Protoplasts aggregate over time and the 1 mL pools were not pipetted for homogeneity. Rather, a fluorescence stereomicroscope was used to manually scan the sample and gather representative images in areas with moderate density of protoplasts. Strong GFP fluorescence was detected in both S and M cells 1 and 2 days after transfection.

**Figure 4.**
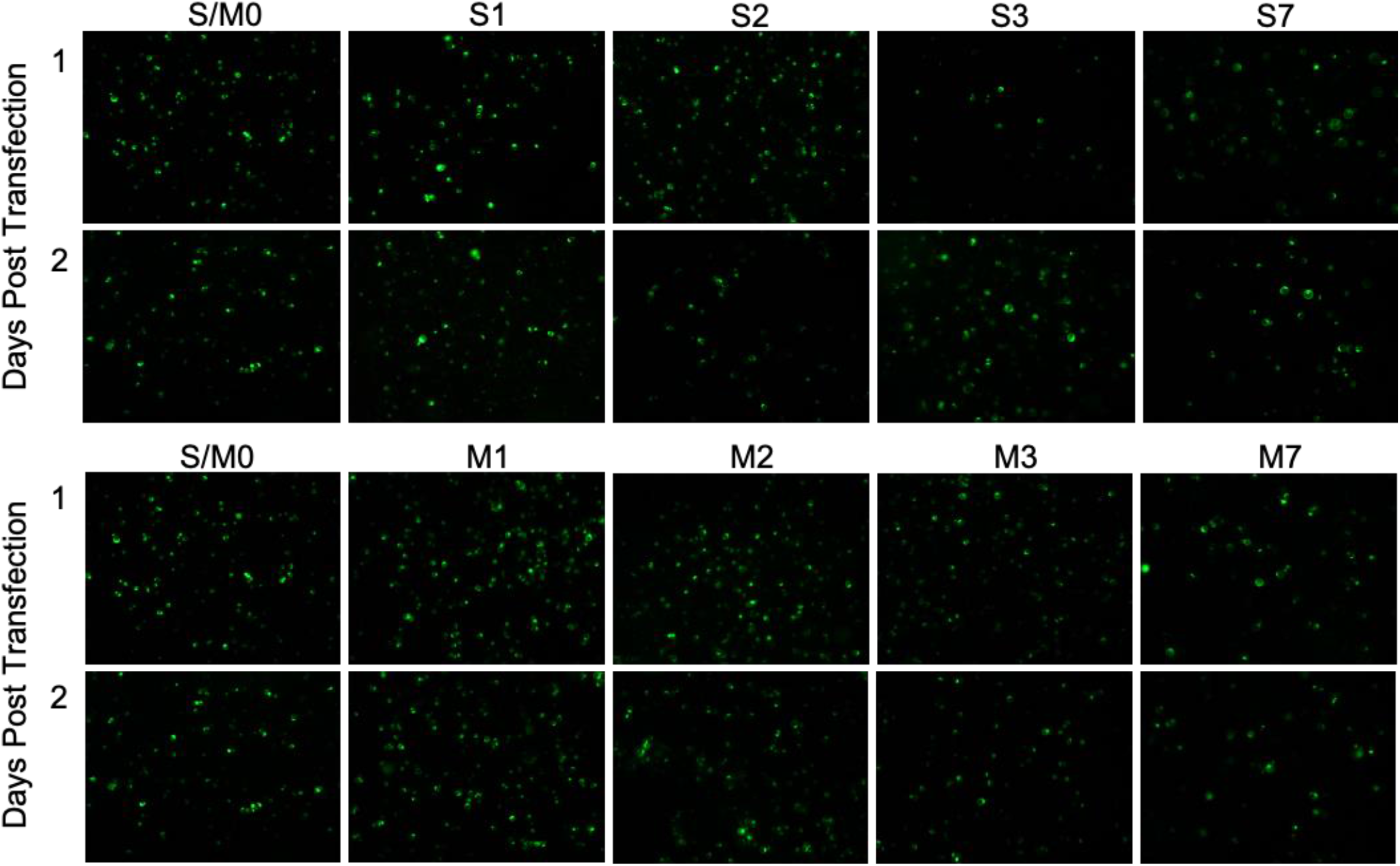
GFP expression in S and M callus-derived protoplasts of different ages. S (top panel) and M (bottom panel) protoplasts of different ages were transfected with pAct1sGFP-1 and imaged for GFP fluorescence. Images were taken at 80X magnification on a Leica M165 fluorescence microscope. Transfected protoplasts were in 1 mL WI solution pools in 12-well culture plates.

Transfection efficiency was calculated 2 days after plasmid transfection as a percentage of GFP positive protoplasts over total live protoplasts, as determined by fluorescence microscopy and Evans Blue staining on a hemocytometer (Fig. 5). For S/M0 protoplasts, transfection was highly efficient, producing 73.5% GFP expressing protoplasts. S1 transfection efficiency was comparable, at 69.5%. Taken together with the results from the previously described viability assay, the data suggest that S1 protoplasts are comparable in value to freshly isolated S/M0 protoplasts. This finding facilitates novel flexibility in research, allowing assays to be performed 24 hours after initiation of the protoplast isolation method with little to no loss of efficacy and data. Though transfection efficiency declines over time for both S and M protoplasts, it does not fall below 15% within 7 days (Fig. 5). Moreover, it is conceivable that certain assays, for example protein localization, do not require optimal transfection efficiency or viability. The data provided here allow for the informed design and versatile scheduling of protoplast experiments with a quantified summary of expected losses of viability and transfection efficiency over time.

**Figure 5.**
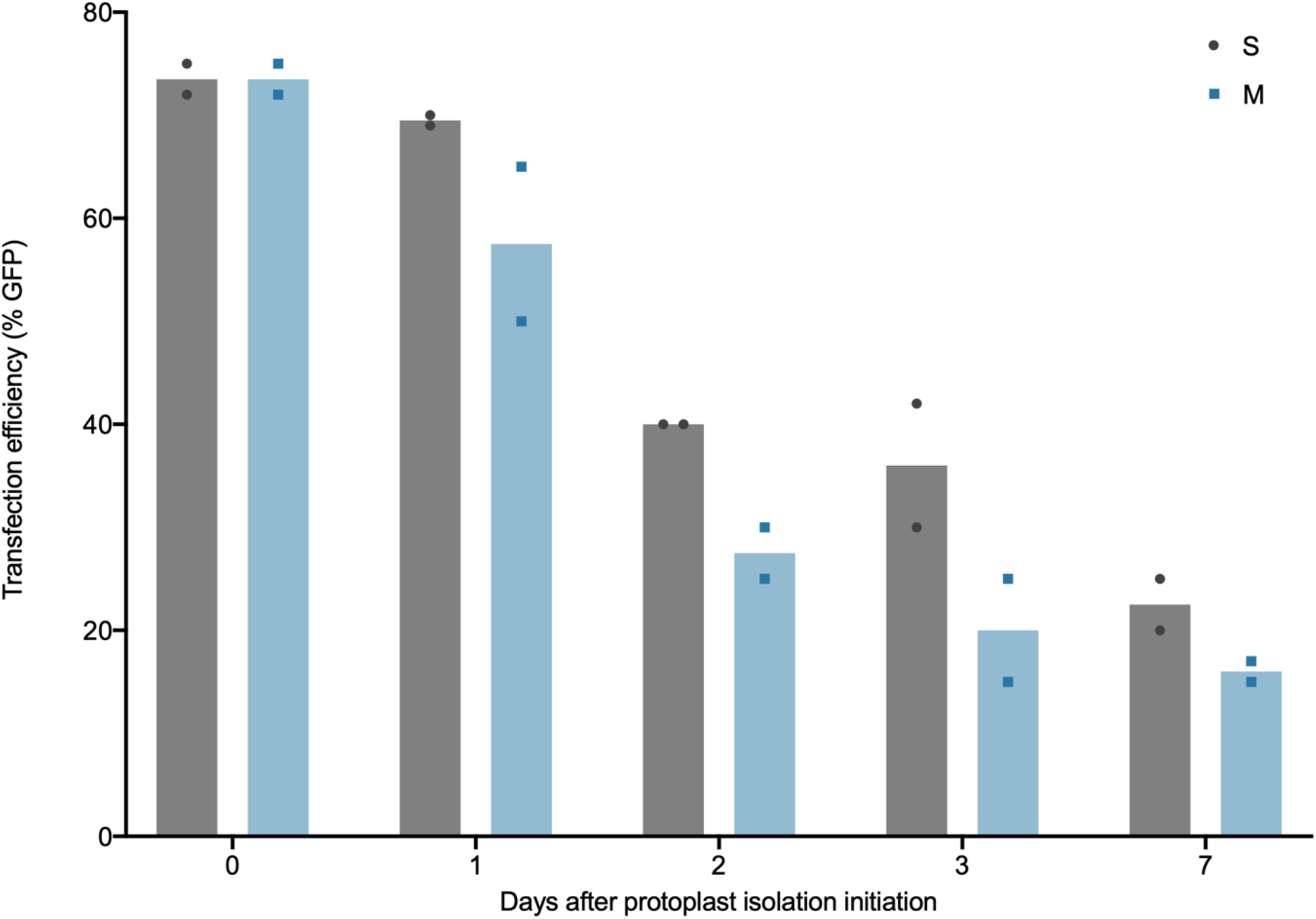
Transfection efficiency in S and M protoplasts isolated from rice calli. The percentage of GFP fluorescence-positive protoplasts were calculated after transfection with pAct1IsGFP-1 to determine plasmid DNA transfection efficiency in S and M callus-derived rice protoplasts of different ages.

### Gene editing via Cas9 ribonucleoprotein transfection

Given the flourishing field of genome editing, it was critical to ensure that this method was suitable for such studies. To demonstrate this, S/M0 as well as S1 and M1 protoplasts were transfected with *in vitro* assembled Cas9-gRNA ribonucleoproteins targeting a single locus (Fig. 6). For comparison with protoplasts isolated via a previously published method [22], protoplasts derived from rice seedlings were also transfected. Editing at the Cas9 cleavage site was identified and quantified through NGS. Editing rates for S/M0 protoplasts and seedling-derived protoplasts were similar (Fig. 6), indicating that the present protocol can be used confidently in genome editing studies.

**Figure 6.**
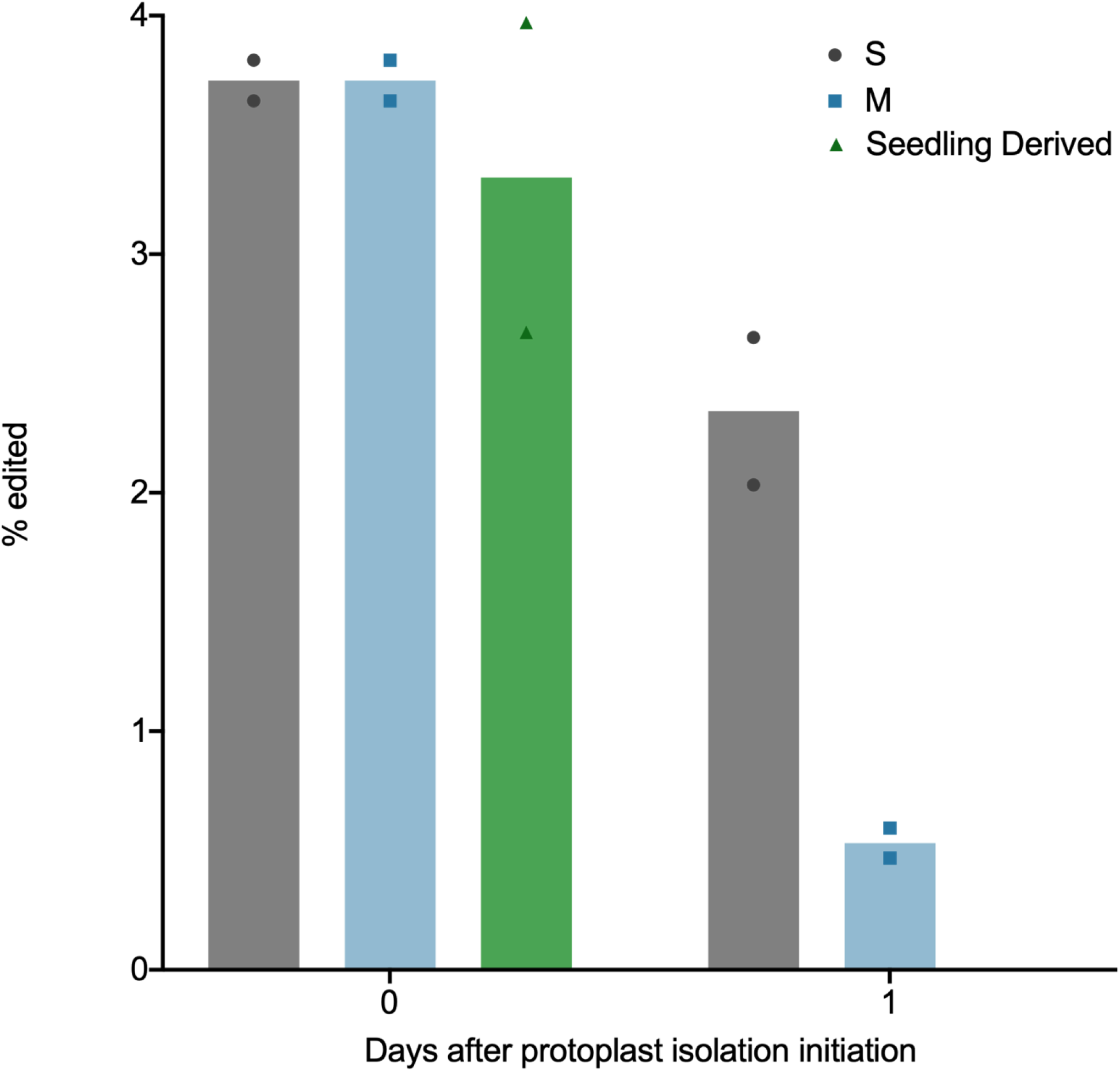
Genome editing efficiency in rice protoplasts isolated from rice calli. Editing efficiency of Cas9 and gRNA ribonucleoproteins in S and M callus-derived rice protoplasts of different ages was compared to seedling-derived rice protoplasts

## Conclusion

The current study describes an embryogenic rice callus-derived protoplast isolation method that avoids the growth of numerous rice seedlings or induction and maintenance of a suspension culture. It also includes optional pause points during and after protoplast isolation. The ability to pause the protocol as well as utilize viable stored M protoplasts increases flexibility in schedules and experimentation for researchers. Because the process of obtaining donor material through isolation of protoplasts and transfection is performed under sterile conditions in its entirety, the protoplasts can be maintained without contamination for time course experiments from transfection through subsequent 7 days or longer. In addition, we demonstrate that the protoplasts produced from this method are competent for transfection of both DNA and RNPs, suitable as transient expression systems, and effective for CRISPR-Cas9 based genome editing assays.

## Abbreviations

2,4-D: 2,4-dichlorophenoxyacetic acid
BAP: 6-benzylaminopurine
MES: 2-(*N*-morpholino)ethanesulfonic acid
PEG: polyethylene glycol
MMG: mannitol magnesium
WI: washing and incubation
CTAB: cetyl trimethylammonium bromide
RNP: ribonucleoprotein
gRNA: guide RNA
SNP: single nucleotide polymorphism
CRISPR: clustered regularly interspaced short palindromic repeats
Cas9: CRISPR-associated protein 9
GFP: green fluorescent protein
NGS: next generation sequencing
rpm: revolutions per minute

## Declarations

### Ethics approval and consent to participate

Not applicable.

### Consent for publication

Not applicable.

### Availability of data and material

All data generated or analyzed during this study are included in this published article.

### Authors’ Contributions

SP and M-JC designed the experiments and wrote the manuscript. SP, JT and M-JC performed the experiments. JHDC, BS and M-JC supervised the experiments. All authors read and approved the final manuscript.

## Acknowledgments

We thank Innovative Genomics Institute (IGI) scientists, Netravathi Krishnappa and Stacia Wyman, for NGS library preparation and sequencing work and for assistance with analysis of sequencing data. Figure 1 was created with Biorender.com.

